# Short-Beaked Echidna (*Tachyglossus aculeatus*) Home Range at Fowlers Gap Arid Zone Research Station, NSW

**DOI:** 10.1101/2020.11.02.364687

**Authors:** Georgia J. Badgery, Jasmin C. Lawes, Keith E.A. Leggett

## Abstract

Echidnas *(Tachyglossus aculeatus)* are found Australian-wide and appear to be remarkably well-adapted to arid zones, yet, nearly all echidna research has been conducted in temperate, tropical and alpine zones. This study investigated the home range and movement of echidnas in western New South Wales. Radio telemetry tracking was used to locate the echidnas daily during the study period (March-May 2019 and August 2019); the home range was 1.47± 1.21 km^2^. This is over twice the reported home range of temperate environments (>0.65 km^2^) suggesting that echidnas exhibit larger home ranges in arid zones. This study provides insight into the movement and home range of echidnas in arid zones, revealing that desert echidnas have large home ranges, dependent on the availability of resources.

## INTRODUCTION

Over the past 200 years, Australian terrestrial mammals have suffered a population decline over 90% (1), additionally, over 30% have become extinct or face possible extinction, primarily due to land clearing and the introduction of exotic species (1, 2). Medium-sized mammals (34-4200g) have been the worst affected, with 25% loss (2, 3). One exception is the short-beaked echidna (*Tachyglossus aculeatus*), which has shown little to no range or population declines despite habitat loss and fragmentation (2). Echidnas and their prey are not restricted by habitat, and they do not require vegetation for shelter, contributing to their success (2). Echidnas are also able to tolerate extreme Australian environmental conditions including drought and fire due to their low metabolic rate (2). Echidnas are likely to be more ecologically important in arid zones compared to zones that receive higher rainfall since they contribute to a larger percentage of the biota (4).

Animals move and migrate for many reasons; to find food, shelter or mates or avoid predation and competition. However, their movements are not random, rather, they are restricted to defined areas, or home ranges (5). This paper defines home range as the area used predictably and regularly for daily metabolic needs (6, 7). Home range research has greatly extended our understanding of the ecology and behaviour of animals (5), and generally involves tracking individuals using a locational device (i.e. GPS or radio-telemetry). Although radio-telemetry involves the observer to physically follow and locate the tagged animal, it is a reliable method of tracking animals that do not have large ranges, it is also significantly cheaper than remote tracking using GPS. The versatility of radio-telemetry tracking means it has become central in studying the ecology of wildlife throughout the world (8, 9). This tracking technique has also been used to investigate the home range and daily movements of a variety of animals globally, including African elephants (*Loxodonta africana*) in Namibia (6),r birds (vultures and condors and the and invertebrates such as the tiger spiketail dragonfly (*Cordulegaster erronea*) (8, 10).

Both Minimum Convex Polygon and Kernel analysis have been widely used for home range analysis (6, 7) despite their marked difference in calculation techniques. The Minimum Convex Polygon technique draws the smallest possibly polygon around all of the points in the home range, ensuring that the interior angles do not exceed 180 degrees. Contrastingly, Kernel analysis uses the density of points to create a statistically weighted home range (11).

Both methods have their strengths and weaknesses, so this study uses both to try to accurately capture the home range of echidnas in an arid zone (7,11).

Echidnas are reported to remain within a defined home range if the habitat is suitable (12). The first recorded home range of echidnas in Australia using radio-telemetry tracking was calculated by taking the furthest distance between two sightings of the same echidna (12). This study was conducted on Kangaroo Island and reported an average home range of 800m in diameter (approx. 0.5 km^2^). The home range and movement ecology of echidnas has since been studied in semi-arid and temperate zones of Western Australia and Tasmania (respectively)(13, 14) as well as the Sub-Alpine Snowy Mountains (15) The studies have used more sophisticated methods of home range calculation such as using minimum convex polygons and kernels. There appears to be some agreement between these home range studies despite the use of different methods, reported that the average home range of echidnas in the Snowy Mountains was 0.42±0.20 km^2^ (15) and 0.50±0.25km^2^ in south-eastern Queensland (6). These studies also demonstrated an overlap of conspecific home ranges and suggested that adult echidnas remain in their home range their entire life.

There is, however, limited information on echidna foraging ecology in arid zones despite these areas making up 70% of Australia’s land area (16). These ecosystems generally have lower biodiversity than higher rainfall areas (4). In arid environments, productivity is greatest in the riparian zones; where aquatic ecosystems transition to terrestrial (17). In arid zones, the riparian ecosystems are often comprised of ephemeral flood out zones, where the surface water is low, unpredictable and irregular (17). Accordingly, fertile areas are concentrated in patches amongst areas of extreme infertility (18).

This study will investigate the home range and daily movement of echidnas in an arid zone and compare this to previous studies of echidna home range in more temperate regions.

Providing an insight into the trends of home range in different climatic zones. We hypothesise that home ranges will be lager in arid environments compared to higher rainfall areas.

## MATERIALS AND METHODS

### Study Site

This study was conducted at Fowlers Gap Arid Zone Research Station (31°05’S, 141°43’E), 110 km north of Broken Hill, NSW (Fig. 1) between March 2018 and September 2019. The station was established in 1966 by the University of New South Wales as both a research and a working sheep station (19). The climate at Fowlers Gap is arid, with a 50-year mean annual rainfall of 230.7 mm, although this is highly variable (19, 20). In summer, daily temperatures exceed 30°C, while the temperatures in winter are mild, rarely falling below 0°C (19).

**Figure 1.**
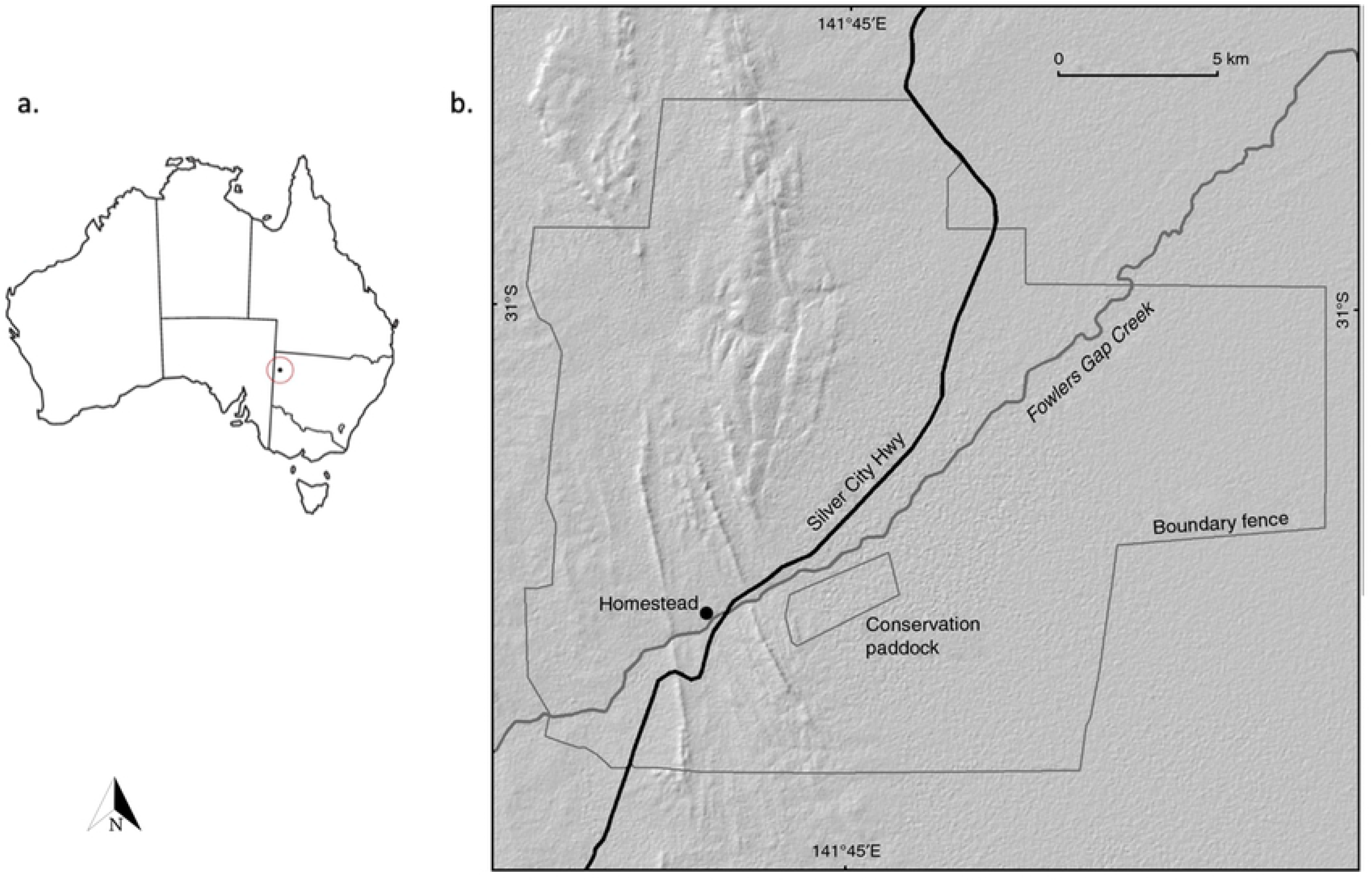
a. Map of Australia indicating position of Fowlers Gap Arid Zone Research Station. b. Inset of Fowlers Gap Arid Zone Research Station.

During and for 2 years prior to this study, Fowlers Gap has been experienced a drought with annual rainfalls of 86.6 mm in 2017, 48.4mm in 2018 and 40.8mm in 2019 (until 30 September) (20). Arid zones are particularly susceptible to climatic changes, and this prolonged drought is likely to have altered the landscape and biodiversity of the region (4).

The study area covered 15 km^2^ around the main homestead, encompassing Fowlers Creek, Homestead Creek and Gum Creek. The area is characterised by arid shrubland, predominately saltbush (*Atriplex vesicaria*) (21, 22), while perennial vegetation is mostly absent. River red gums (*Eucalyptus camaldulensis*) and prickly wattle (*Acacia victoriae*) dominate the riverine woodlands, providing ample shelter for echidnas (21, 22). The focus of this study was the riparian vegetation zone and areas surround the large artificial lakes used for stock and domestic water supply. The ephemeral creek habitats are characterised by steep, well defined banks surrounding narrow flow beds as well as and sparse trees both in the flow bed and along the bank shelves. The artificial lake habitat consists of a semi-permanent water hole surrounded by indistinct banks, ephemeral flow bed and flood out. The bank and ephemeral flood out area is covered with river red gum and a dense layer of leaf litter which creates a cool, moist environment (19).

### Home Range Analyses

Tx-VHF Transmitters (Model F1840) were fitted to ten echidnas in 2018 and three echidnas were tagged during the study, in April 2019. These transmitters were manufactured by Advanced Telemetry Systems. They have a battery life of 787 days and send 40 pulses (between 150-152MHz) per minute (ppm).

Echidnas were found on foot by direct searching throughout the study area (around the Folwers Gap Homestead) while the animals were foraging or moving. Once echidnas were located, spines on the acnestis were clipped to ~5 mm long, and a 20 g transmitter was attached using a fast-setting non-toxic epoxy resin (Gorilla Glue Inc. USA). The transmitters did not exceed 5% of the echidna’s body mass, which is the recommended limit (23). Each echidna was weighed upon capture and categorised as either juvenile (<3 kg) or adult (>3 kg) (24, 25). Echidnas were identified as females if there was an absence of hind-leg spurs (23). However, other sex-specific characteristics such as pouch development in females, or penis protrusion in males were not reliably identified (13, 24).

Over a 2-year period, thirteen adult and sub-adult echidnas were tracked to calculate home range of echidnas at Fowlers Gap. The echidnas were numbered: E1-13, with number 1 being the earliest tagged, on 16 March 2018 and number 13 being tagged last, on 27 April 2019. Of these, E1, E2, E4, E5, E6 and E13 were reliably be identified as females, based on the absence of hind leg spurs (22). Weights of echidnas, at the time of capture, ranged from 1.1-4.8kg. E7, E8, E12 and E13 were excluded from home range analyses since they were observed less than 15 times (6). No signal was located for E9 and E12 during September, 2019. E1 and E4 were found dead and buried on 29/11/18, E7 was found dead on 22/2/19. E13 lost her radio tracker sometime between May and August 2019. The largest (4.8kg) and all tagged young animals (<1.2kg) (24), were among those confirmed dead.

Echidnas were located daily using a VHR receiver (ATS R2000) and a handheld 3 Element Folding Yagi Antenna (ATS). Searching commenced on hills or creek banks near the last recorded location of the animal and continued until reliably located the echidna by either: physically observing the echidna or finding a pin-pointed signal above a burrow. Once located, the foraging or resting site was marked with yellow flagging tape and the geographical coordinates were obtained using a Garmin handheld GPS with at least 4 m accuracy.

Similarly with a previous home range studies (6), independent locations were determined by:

1. All locations of echidnas not in shelters, or
2. All locations of echidnas in shelters, unless they have previously occupied that shelter with no evidence that they have left the shelter and then returned.

Augee et al. (15) suggested that at least 15 data points are required to calculate home range reliably. Hence, animals with less than 15 independent data points were not used for home range analysis.

### Statistical Analyses

Home range of each echidna was mapped using R and Google Earth Pro. Home range of the echidnas was estimated by minimum convex polygon (MCP) analysis (95%) using the *Minimum Bounding Geometry* (6, 7). The Kernel function in R was used to further analyse home range. The fixed kernels used during this analysis were 50% and 95% of loci, which correspond to ‘core area’ and ‘peripheral area.’ The smoothing factor of the Kernel was adjusted so that the area of the 95% kernel was approximately equal to the area of the MCP, although this was not possible when echidnas appeared to leave their home range (11).

## RESULTS

Table 1 summarises the data collected from the nine echidnas that were used for home range analysis including capture weight and whether the animal was identified as female. The home range of echidnas at Fowlers Gap (95% MCP) ranged between 0.02 km^2^ to 3.65km^2^. The peripheral areas (95% Kernel) ranged between 0.31 km^2^ to 1.87 km^2^ while the core areas (50% Kernel) were between 0.04 km^2^ to 0.38 km^2^ (Table 1). The 95% Kernel was much smaller than the 95% MCP for echidnas E4 and E6. However, the 95% Kernel was much larger than the 95% MCP for E5. Seven out of nine of the echidnas shared a home range with at least 2 other echidnas.

**Table 1.** Home ranges of echidnas determined by three different calculation methods: 95% Kernels (peripheral area), 50% Kernels (core area) and 95% Minimum Convex Polygons (MCP). “F” indicates that the echidna was positively identified as female. SD: Standard deviation.

| Echidna ID | Weight on capture (kg) | Number of independent locations | Kernel (km<sup>2</sup>) |  | MCP (km<sup>2</sup>) | Number of conspecifics within the home range |
| --- | --- | --- | --- | --- | --- | --- |
|  |  |  | 50% | 95% | 95% |  |
| E1 | 1.24 F | 19 | 0.16 | 0.79 | 0.68 | 0 |
| E2 | 2.9 F | 41 | 0.12 | 0.54 | 0.44 | 2 |
| E3 | 4.02 | 73 | 0.23 | 1.42 | 1.20 | 5 |
| E4 | 2.9 F | 16 | 0.08 | 0.50 | 3.56 | 0 |
| E5 | 1.1 F | 32 | 0.04 | 0.31 | 0.02 | 2 |
| E6 | 2.1 F | 90 | 0.38 | 1.87 | 3.38 | 6 |
| E9 | 2.8 | 59 | 0.17 | 0.93 | 0.79 | 6 |
| E10 | 2.3 | 59 | 0.13 | 0.61 | 0.36 | 2 |
| E11 | 3.5 | 21 | 0.24 | 1.05 | 1.01 | 6 |
| Mean |  |  | 0.17 | 0.89 | 1.27 |  |
| SD |  |  | 0.10 | 0.49 | 1.29 |  |

Figure 2 shows the difference between MCP and Kernel analyses using the E6 as an example. The 95% kernel is fragmented, displaying three distinct clusters that extend beyond the individual data points which reflects where an animal spends 95% of its time. The MCP creates one distinct shape that encompasses all of the individual data points.

**Figure 2.**
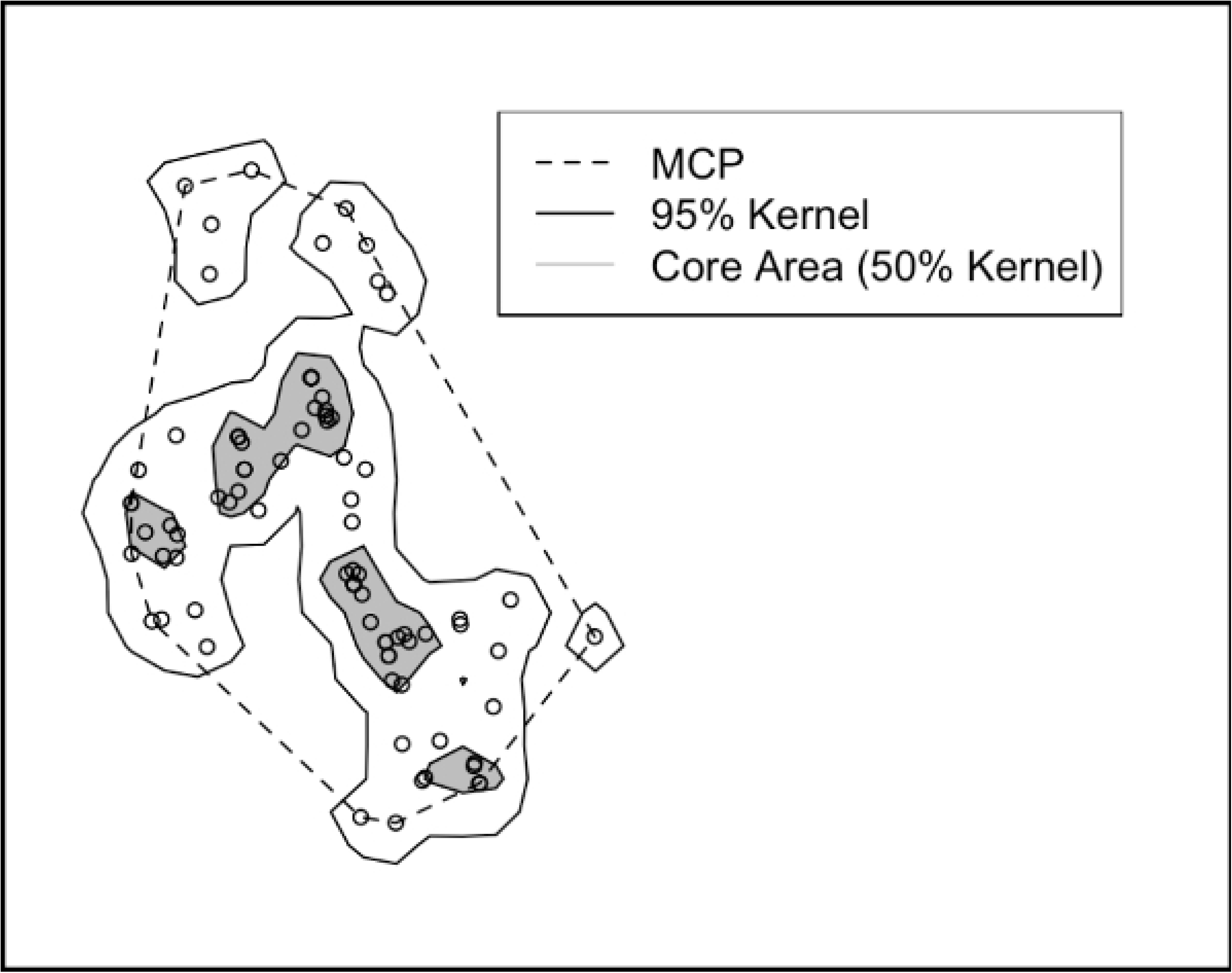
Minimum Convex Polygon, 95% Kernel and Core Area of E6 including individual data points.

Figure 3 maps the MCP of eight echidnas (E1, E2, E3, E4, E6, E9, E10 and E11). E5 was excluded because the MCP was too small to be clearly included on the map. E2, E3, E6, E9 E10 and E11 are clearly clustered around the Fowlers Gap Homestead while E1 and E4 spread out to the north and south (respectively) of the homestead.

**Figure 3.**
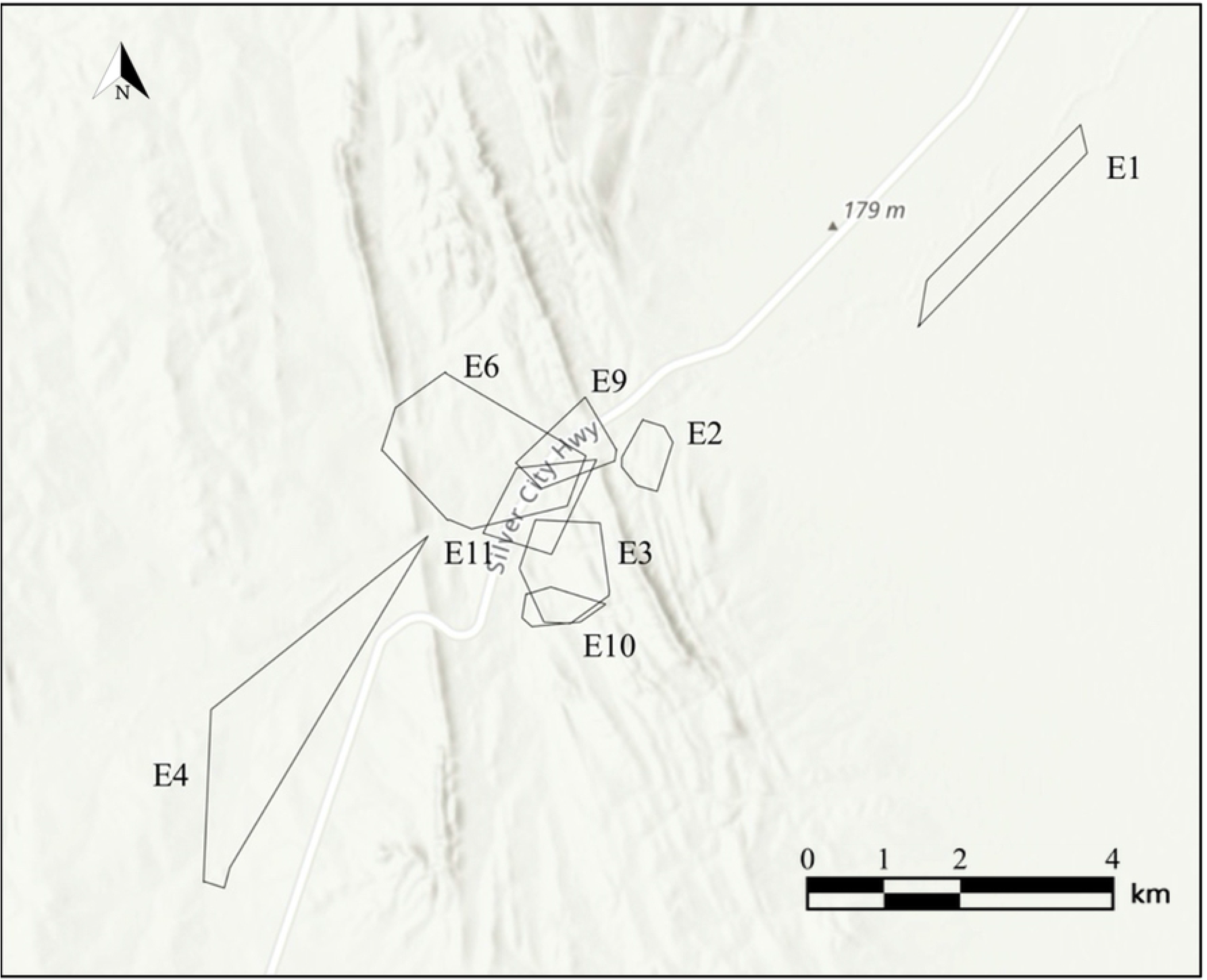
Home range of echidnas E1, E2, E3, E4, E6, E9, E10 and E11 using MCP.

Table 2 shows the percent overlap between the home ranges of two echidnas. The percentage overlap between home ranges was found to vary between zero and 84.2% (mean = 6.61%, SD = 19.8%). E5 has a small home range which is 100% included in the home ranges of E6 and E11, while E5 only makes up 2 and 10 percent of E6 and E11’s home range, respectively.

**Table 2.**
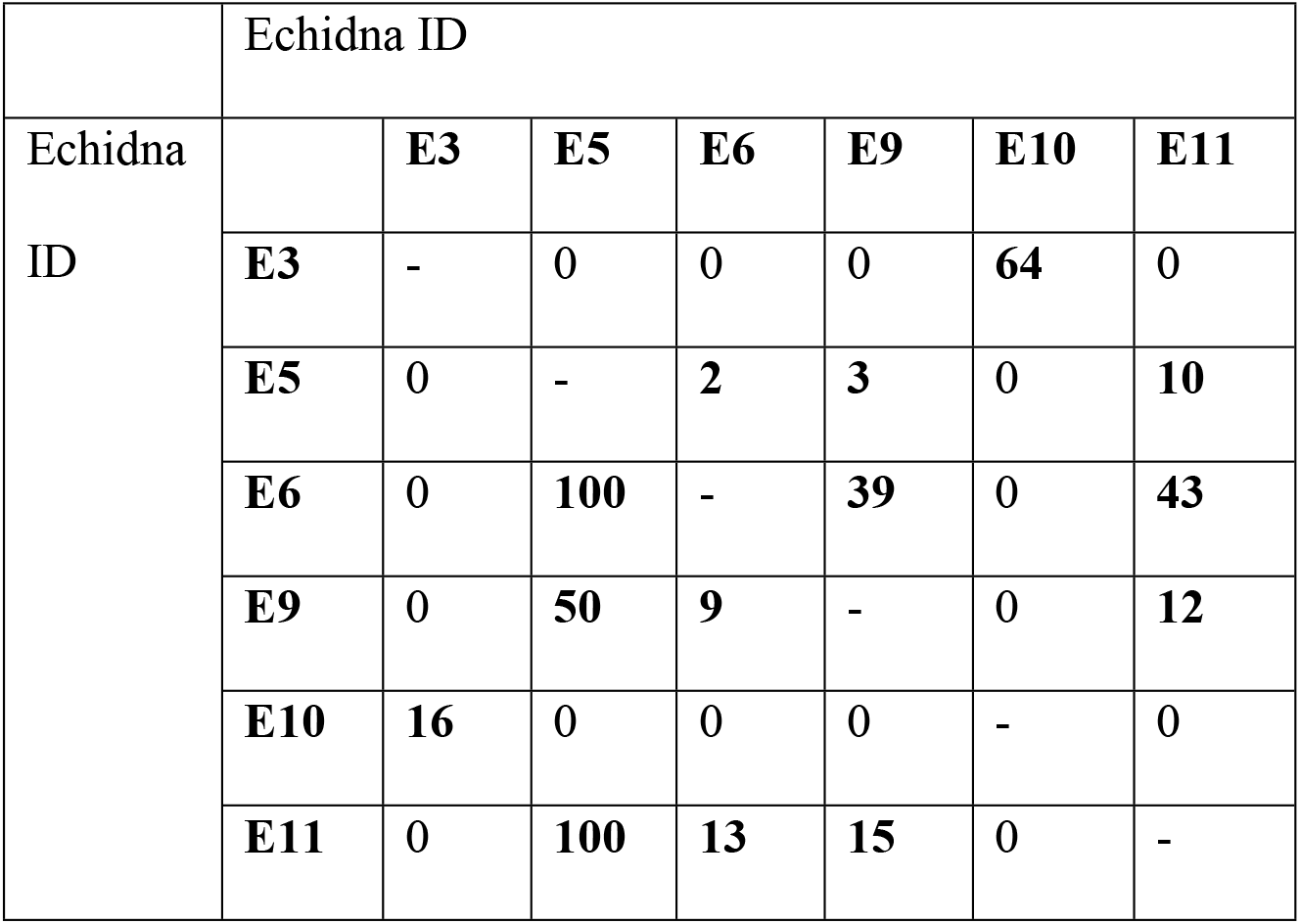
Percentage overlap of echidna home ranges

|  | Echidna ID |  |  |  |  |  |  |
| --- | --- | --- | --- | --- | --- | --- | --- |
| Echidna ID |  | **E3** | **E5** | **E6** | **E9** | **E10** | **E11** |
|  | **E3** | - | 0 | 0 | 0 | **64** | 0 |
|  | **E5** | 0 | - | **2** | **3** | 0 | **10** |
|  | **E6** | 0 | **100** | - | **39** | 0 | **43** |
|  | **E9** | 0 | **50** | **9** | - | 0 | **12** |
|  | **E10** | **16** | 0 | 0 | 0 | - | 0 |
|  | **E11** | 0 | **100** | **13** | **15** | 0 | - |

## DISCUSSION

This paper shows that the home range echidnas at Fowlers Gap was 1.15±0.60 km^2^. In the context of previous research that this is larger than has been observed anywhere else in Australia, being approximately double the home range of echidnas in the semi-arid Western Australian Wheatbelt (2) and over double that in temperate, tropical, Mediterranean and sub-alpine zones (summarised in Table 3.). The larger home range is likely due to the low-quality habitat provided across the arid zone in contrast to areas that receive more rainfall, including semi-arid zones (4). Home range was concentrated around riparian zones, likely due to the larger amounts of vegetation, and hence, shelter and prey availability (26). Arid zones have fewer and more scattered resources than higher rainfall areas (4). Since the relationship between energy needs and food availability are known to contribute to home range size, it is likely that at Fowlers Gap echidnas needed to utilise larger home ranges to meet their energy requirements (27, 28).

**Table 3.**
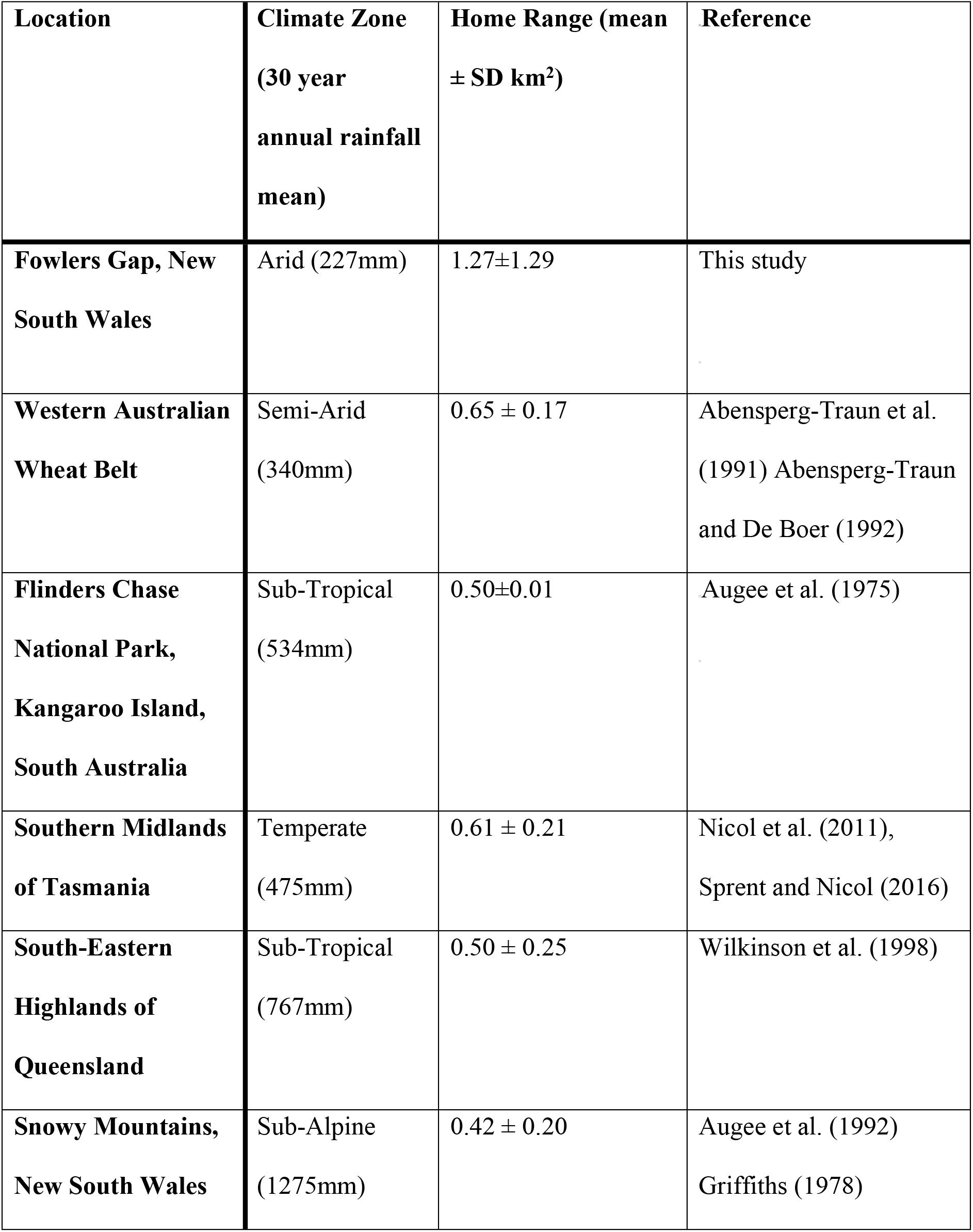
Comparison of the home range size of echidnas within different climate zones.

| **Location** | **Climate Zone**<br>**(30 year**<br>**annual rainfall**<br>**mean)** | **Home Range (mean**<br>**± SD km<sup>2</sup>)** | **Reference** |
| --- | --- | --- | --- |
| **Fowlers Gap, New**<br>**South Wales** | Arid (227mm) | 1.27±1.29 | This study |
| **Western Australian**<br>**Wheat Belt** | Semi-Arid<br>(340mm) | 0.65 ± 0.17 | Abensperg-Traun et al.<br>(1991) Abensperg-Traun<br>and De Boer (1992) |
| **Flinders Chase**<br>**National Park,**<br>**Kangaroo Island,**<br>**South Australia** | Sub-Tropical<br>(534mm) | 0.50±0.01 | Augee et al. (1975) |
| **Southern Midlands**<br>**of Tasmania** | Temperate<br>(475mm) | 0.61 ± 0.21 | Nicol et al. (2011),<br>Sprent and Nicol (2016) |
| **South-Eastern**<br>**Highlands of**<br>**Queensland** | Sub-Tropical<br>(767mm) | 0.50 ± 0.25 | Wilkinson et al. (1998) |
| **Snowy Mountains,**<br>**New South Wales** | Sub-Alpine<br>(1275mm) | 0.42 ± 0.20 | Augee et al. (1992)<br>Griffiths (1978) |

| Location | Climate Zone<br>(30 year<br>annual rainfall<br>mean) | Home Range (mean<br>± SD $\text{km}^2$) | Reference |
| --- | --- | --- | --- |
| Fowlers Gap, New<br>South Wales | Arid (227mm) | $1.27 \pm 1.29$ | This study |
| Western Australian<br>Wheat Belt | Semi-Arid<br>(340mm) | $0.65 \pm 0.17$ | Abensperg-Traun et al.<br>(1991) Abensperg-Traun<br>and De Boer (1992) |
| **Flinders Chase National Park, Kangaroo Island, South Australia** | Sub-Tropical<br>(534mm) | 0.50±0.01 | Augee et al. (1975) |
| **Southern Midlands of Tasmania** | Temperate<br>(475mm) | 0.61 ± 0.21 | Nicol et al. (2011),<br>Sprent and Nicol (2016) |
| **South-Eastern Highlands of Queensland** | Sub-Tropical<br>(767mm) | 0.50 ± 0.25 | Wilkinson et al. (1998) |
| **Snowy Mountains, New South Wales** | Sub-Alpine<br>(1275mm) | 0.42 ± 0.20 | Augee et al. (1992)<br>Griffiths (1978) |

Echidnas are thought to have a stable range that they occupy throughout their lives (12) however, during this study, three adult echidnas could not be found after being tagged in the study area. It is assumed that these echidnas dispersed from the study area, as no signal or carcasses were found after extensive searching. This movement could be evidence of a dispersal event. Dispersals are most commonly seen in young males (29), but since these were absent from this study, it is more likely that the movement out of the area was in response to current drought conditions (2). Drought conditions may have caused previous home ranges to become too resource-poor, forcing animals to seek out new areas with more available resources. Hence, it appears that the resource availability could be a factor in determining echidna home range size and movements. One echidna, E5, appeared to leave its home range and was not able to be located after May 2018, this could account for the disparity between home range recorded by the 95% MCP (0.02 km^2^) and the 95% Kernel (0.31 km^2^). E5, was the lightest (youngest) echidna and had the smallest home range. It also had the highest degree of overlap with two echidnas sharing 100% of its home range and another sharing 50%.

The home ranges of echidnas at Fowlers Gap varied between individuals. E1, E2 and E10 that displayed a home range similar in size to that of sub-tropical and sub-alpine zones where resources are abundant (4, 6, 15)(Table 3). The home range of E10 was concentrated around an artificial lake suggesting that this area may be more resource-rich compared to the creeks and non-riparian zones. Hence, the lake habitat appears to provide a superior habitat to that of the creek beds and non-riparian zones. Non-riparian zones are reported to be more resource-poor than riparian zones (18), this could explain why E10 had a smaller home range than echidnas in a resource poor area. However, it is not why the home range around creeks is larger than around artificial lakes. It is possible that the artificial lakes and watering points may support a higher carbon concentration (more trees, leaf litter and detritus), and hence be more resource-rich than the natural ephemeral creeks (18).

E4 and E6 exhibited very large home ranges (Table 1). E4 appeared to disperse and leave its home range before being found dead in November 2018, this likely accounts for the large home range calculation. This dispersal was possibly due to a lack of resources forcing it to unsuccessfully seek out a new home range. E6 however, consistently utilised its home range for the entire duration of this study. This echidna was mostly found along two creek lines and within two gardens. 43% of E11’s home range overlaps with E6 overlapped, sharing a resource rich area around the homestead and along a creek. However, E11 demonstrated a significantly smaller home range than E6. It is unclear as to why E6 used such a large home range when it shared a resource rich area with E11. This study showed that echidnas had a smaller degree of overlap of home ranges than in previous studies, an average overlap of 6.61% compared to 24% in south-eastern Queensland and extensive overlap in the Snowy Mountains (6, 15).

As arid zones have more sparse recourses and lower biodiversity than higher rainfall zones (4), it is likely that a particular home range will not be able to sustain multiple echidnas, or that echidna home ranges are more strongly linked to resource availability than their temperate counterparts. Hence, it is likely that echidnas in arid zones require larger home ranges in order to meet their energy requirement. The low resource availability could also force some echidnas to disperse and seek out alternate foraging areas. It is possible that E5 exhibited this behaviour during this study by initially foraging and occupying 100% of its home range with 2 echidnas, then moving to find alternate resources. It is also possible that E5 was a juvenile (being the smallest echidna at 1.1kg upon capture) and that this dispersal was due to it being weaned from its mother.

This study has provided strong evidence to suggest that Echidna home range is much larger in arid zones compared to semi-arid, temperate, subtropical and sub-alpine zones. Further research into the driving factors of echidna home range size across climatic zones could be useful in creating a model and hence predicting the requirements for the success of the species.

## ACKNOWLEDGEMENTS

This study was undertaken on research permit (Approval number: 18/3B) issued by the UNSW Animal Ethics Committee, and NSW National Parks and Wildlife Service scientific license number: SL102050.

We also thank Kasin Ki, Corrine Edwards, James Badgery, Michael Letnic, Zara Badgery, Michelle Ayoub and The Dowling Family for assistance in the field. Additionally, we would like to thank Elizabeth Weir for reading and editing the text.

